# Thyroid Hormones-Thyrotropin Interrelationships in Thyroid Dysfunction States

**DOI:** 10.1101/2020.06.26.173252

**Authors:** Tika Ram Lamichhane, Sandeep Prashad Pant, Binod Lamichhane, Sharma Paudel, Binod Kumar Yadav, Hari Prasad Lamichhane

## Abstract

Objective of this research is to explore the complex correlations among FT3, FT4 and TSH in thyroid dysfunction states. A total of 3425 outpatients were included to analyze their thyroid function tests measured at the university teaching hospital. The subjects under study were categorized as euthyroid (70.45%), subclinical hypothyroid (18.95%), overt hypothyroid (3.30%), subclinical hyperthyroid (5.11%) and overt hyperthyroid (2.19%) patients. The linear and non-linear relations among FT3, FT4 and TSH were modeled statistically to understand their role in thyroid functions. The prevalence of hypothyroidism is higher than hyperthyroidism in which females are more vulnerable than males. The correlation between FT3 and FT4 increases from euthyroidism (r = 0.1, P <0.001) to overt thyroid dysfunctions (r = 0.91, P <0.001). The non-linear complex correlations between FT4 and TSH are significantly explained by well fitted sigmoid curves of four-parameter logistic (4PL) model (R^2^ = 0.97 and P <0.001) in both hypothyroidism and hyperthyroidism interfaced with euthyroidism. The 4PL curves are response trajectories of pituitary TSH which exhibit the turning points of thyroid dysfunction states by the deviation in serum FT4. The negative slopes and correlations observed in the linear fits in TSH *vs* FT3 or FT4 are the indicators of negative feedback mechanism in the thyroid cycle.

## 1. Introduction

Age, gender, ethnicity and geography based study is necessary to evaluate thyroid functions. The reference levels of serum free thyroid hormones (FT3, FT4) and thyroid stimulating hormone (TSH) vary between different laboratories which make remarkable changes in judging thyroid status of the patients.^1-5^ The analysis of thyroid function tests (TFTs) results the complex correlations among the covariates: FT3, FT4 and TSH under the influence of many physiological factors. The hormone relationship is modulated by thyroid and non-thyroid illness, medication, body-mass index, age, gender and antibody status.^1,3,6,7^ Developing from simple linear fits among the TFT values, log-linear relationships between TSH and FT4 came gradually into practice.^1,8,9^ In addition to the log-linear transformation, non-linear relationships between TSH and FT4 have also been discussed based on different models such as sigmoid functions, error function and fourth-order polynomial.^2,10-12^ According to Reichlin *et al*. (1967), TSH secretion is finely regulated for a wide range of FT4 concentrations and there is a curvilinear relationship between TSH and FT4 supporting the negative feedback system of HPT axis.^13^ Midgley *et al*. (2013) suggested for analyzing separately euthyroid, hypothyroid and hyperthyroid panels to understand the correlation between TSH and FT4 because the functional segments appear differently regulated as the trends between these states change.^14^ Hadlow *et al*. (2013) has modeled the clinical data to develop the sigmoid relations between TSH and FT4 in two segments of FT4 <12 pmol/L and FT4 >12 pmol/L. However, their significance in thyroid dysfunction states are undefined.^2^ The correlations among FT3, FT4 and TSH change in the different states of thyroid disorders for which we analyze the big data of the hospital based subjects. In this research, four-parameter logistic (4PL) model has been better applicable for the complex correlations between TSH and FT4 when thyroid disorder grows from euthyroidism to subclinical and overt thyroid dysfunction. The 4PL model explains about Hill’s slope, inflection point and asymptotic values that yield a relation for response trajectory of pituitary TSH by the deviation in serum FT4.

## 2. Materials and Methods

The TFT reports of outpatients were obtained from the laboratory database of Department of Biochemistry, Tribhuvan University teaching Hospital (TUTH), Kathmandu, Nepal. The TFTs were measured by enhanced chemiluminescence immunoassay technique using Vitros 3600 machine form September 2017 to December 2018. A total of 3425 subjects aged 0.50 to 93 years were included in this study. The reference ranges of FT3, FT4 and TSH were 4.26-8.10 pmol/L, 10.20-28.20 pmol/L and 0.46-4.68 mIU/L, respectively. The ethical approval was provided by Institutional Review Board (IRB), TUTH. The data having analytical interference by chronic non-thyroidal illness, impacts of medication, factors confounding the HPT axis response and lack of information such as unknown about entry date, age, gender and missing any one of the TFTs were excluded from this study.

The TFT dataset was categorized into euthyroid, subclinical hypothyroid, overt hypothyroid, subclinical hyperthyroid and overt hyperthyroid subsets. Average TFTs with permissible errors were calculated for each dysfunction state. The data subsets were statistically analyzed with linear fits between FT3 and FT4 and with log-linear fits between ln(TSH) and FT3 or FT4. Also, non-linear complex correlations between FT4 and TSH were analyzed by using 4PL regressions separately in hypothyroid and hyperthyroid states on the basis of euthyroid state. The statistical analyses including linear as well as non-linear fits were carried out using MS Excel-07 and Origin-2017 (Version 94E, OriginLab Corporation, USA) software packages. The results obtained with P-value less than 5% were considered to be statistically significant.

As suggested by Bortolotto *et al*. (2015)^15^ and Azadeh *et al*. (2018)^16^ for the ligand binding assays, the sigmoid equation of 4PL model that supports the non-linear relationship between FT4 and TSH is given by

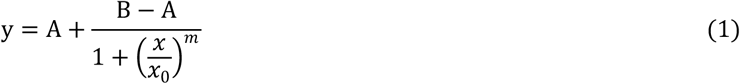

where *y* is ln(TSH), *x* is FT4, A is minimum asymptotic y-value or response value for the maximum standard FT4, B is maximum asymptotic y-value or response value for the minimum FT4, *x*_0_ is inflection point or *x* -value corresponding to half-way between A and B, and *m* is Hill’s slope or steepness of the curve. The 4PL curve is sigmoid symmetric in nature.

## 3. Results and Discussion

In a total of 3425 subjects, there are 70.45% (665M+1748F) clinically euthyroid, 18.95% (164M+485F) subclinical hypothyroid, 3.30% (29M+84F) overt hypothyroid, 5.11% (46M+129F) subclinical hyperthyroid and 2.19% (17M+58F) overt hyperthyroid patients. The patients are more suffering from hypothyroidism than hyperthyroidism in which females (F) are more vulnerable than males (M) as reported in the previous studies.^7,17-20^ The descriptive statistics (mean, standard deviation (SD), minimum and maximum values) of age, TSH, FT4 and FT3 for each of the data sets are depicted in Table 1. The results are statistically significant (P <0.001) while comparing those groups of thyroid states.

**Table 1.**
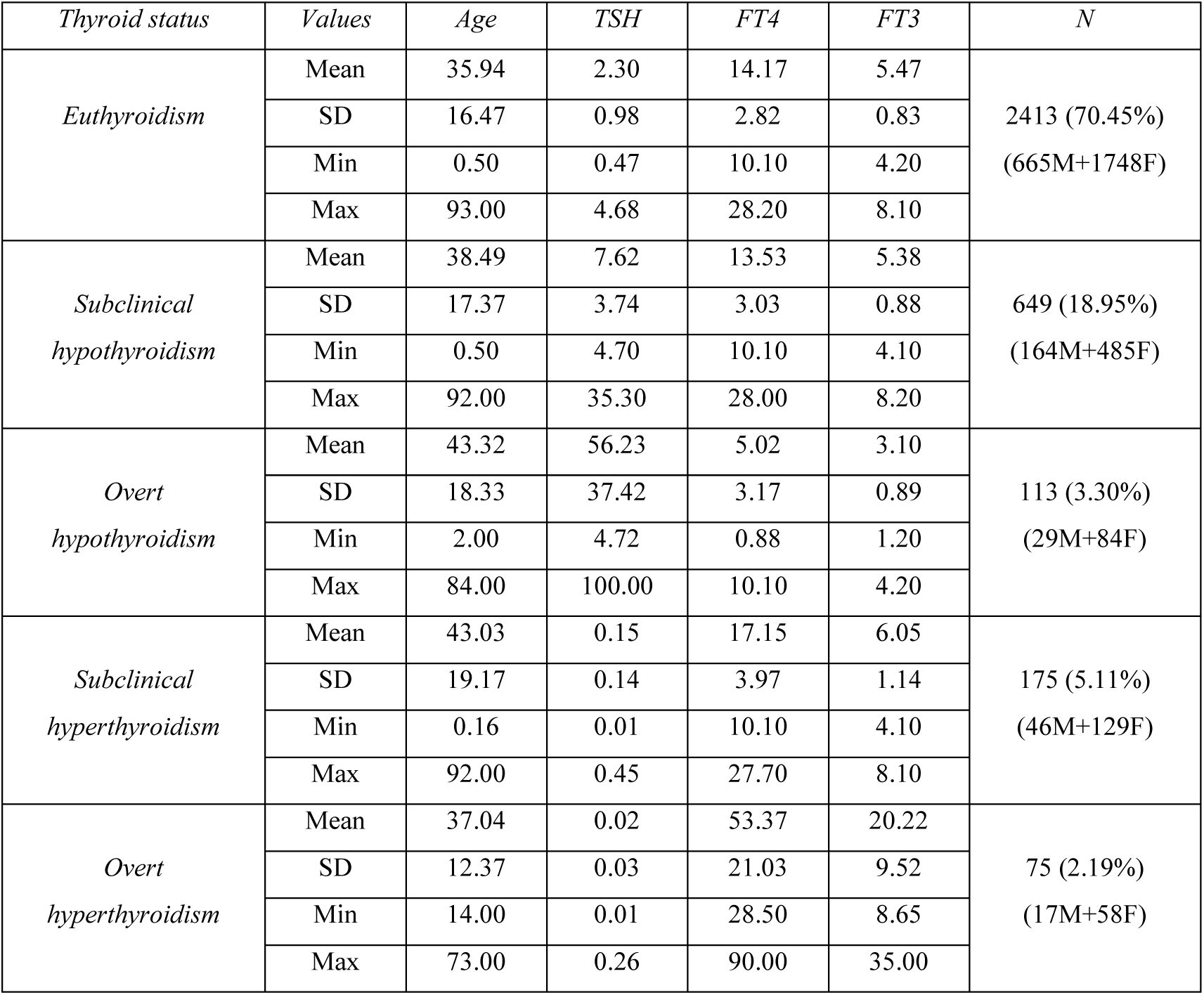
Descriptive statistics of age(year), TSH(mIU/L), FT4(pmol/L) and FT3(pmol/L) in normal and abnormal thyroid conditions of 3425 patients where ANOVA single factor of each TFT has P <0.001 over the different groups

The mean±SD of TSH, FT4 and FT3 are 2.30±0.98 mIU/L, 14.17±2.82 pmol/L and 5.47±0.83 pmol/L, respectively in the euthyroid patients aged 35.94±16.47 years. In the subclinical hypothyroid patients aged 38.49±16.37 years, the values of TSH, FT4 and FT3 are found to be 7.62±3.74 mIU/L, 13.53±3.03 pmol/L and 5.38±0.88 pmol/L, respectively. The overt hypothyroidism appears in the higher age of 43.32±18.33 years in which the elevated TSH is 56.23±37.42 mIU/L whereas the suppressed FT4 and FT3 are 5.02±3.17 pmol/L and 3.10±34.20 pmol/L. The subclinical hyperthyroid patients aged 43.03±19.17 years have the suppressed TSH of 0.15±0.14 mIU/L with the normal FT4 of 17.15±3.97 pmol/L and FT3 of 6.04±1.14 pmol/L. The overt hyperthyroid patients aged 37.04±12.37 (range: 14 to 73) years have highly suppressed TSH of 0.02±0.03 mIU/L with the elevated FT4 of 53.37±21.03 pmol/L and FT3 of 20.22±9.52 pmol/L. Each of the TFTs has P <0.001 over the different groups (Table 1). Our observations also show that the occurrance of overt thyroid dysfunctions is lower in early childhood than in elderly as explained in the earlier studies.^21-23^ The thyroid related researches show that most of the children suffer from congenital thyroid dysfunction and adults suffer from autoimmune thyroiditis, cardiac failure by thyrotoxicosis, hypertension and thyroid nodules or goitre. ^24^

The linear correlation between FT3 and FT4 is stronger in both overt hypothyroidism (r = 0.66, P <0.001) and overt hyperthyroidism (r = 0.91, P <0.001) than in subclinical hypothyroidism (r = 0.21, P <0.001), subclinical hyperthyroidism (r = 0.10, P <0.17) and euthyroidism (r = 0.10, p <0.001) as shown in Fig. 1. The slope of linear fit in FT3 *vs* FT4 also increases from euthyroidism to hypothyroidism and to hyperthyroidism as depicted in Table 2. The increasing slope means the increasing rate of monodeiodination of FT4 to FT3 under the deregulated TSH. In the overt thyroid dysfunction, the extreme values of TSH are taken to be 0.01 and 100 mIU/L. We found significant linear correlations (r = -0.75, P <0.001 in ln(TSH) *vs* FT4 and r = -0.52, P <0.06 in ln(TSH) *vs* FT3) in case of overt hypothyroidism. However, the linear fits in ln(TSH) *vs* FT4 suggest very weak correlations in other cases with P-value ranging from <0.001 to 0.72 as presented in Table 2. Then negative slope and correlation coefficient indicate the negative feedback mechanism of thyroid hormones into hypothalamus and pituitary. This result emphasizes for the non-linear relationship between TSH and FT4 (or FT3). The underlying factors controlling the expression of body function develop the non-linear relationship between TSH and FT4. The related contributing factors are age, sex, body-mass index, HPT-axis function, thyroid and non-thyroid illness, genetics, environmental factors, depression, anxiety and medications.^1,25-27^ The TSH-centered TFT reflects thyroid status in different pathophysiological and physiological conditions.

**Table 2.** Coefficients of linear fit: y = Ax + B among the variables log_e_ of TSH (mIU/L), FT3 (pmol/L) and FT4 (pmol/L) in euthyroidism, hypothyroidism and hyperthyroidism. The included errrors are standard errors in slope and intercept, and the P-value for F-test is obtained from the linear regression analysis

| Thyroid status | Values | $\ln(TSH)$ vs FT4 | $\ln(TSH)$ vs FT3 | FT3 vs FT4 |
| --- | --- | --- | --- | --- |
| Euthyroidism<br>(N = 2413) | Slope (A) | -0.02±0.003 | -0.02±0.01 | 0.03±0.006 |
|  | Intercept (B) | 0.99±0.05 | 0.84±0.06 | 5.00±0.08 |
|  | Pearson's r | -0.11 | -0.03 | 0.11 |
|  | P-value | <0.001 | 0.07 | <0.001 |
| Subclinical<br>hypothyroidism<br>(N = 649) | Slope (A) | -0.005±0.004 | -0.04±0.02 | 0.06±0.01 |
|  | Intercept (B) | 2.02±0.06 | 2.16±0.08 | 4.56±0.15 |
|  | Pearson's r | -0.04 | -0.10 | 0.21 |
|  | P-value | 0.26 | 0.013 | <0.001 |
| Overt<br>hypothyroidism<br>(N = 113) | Slope (A) | -0.23±0.02 | -0.56±0.09 | 0.18±0.02 |
|  | Intercept (B) | 4.84±0.11 | 5.40±0.29 | 2.17±0.12 |
|  | Pearson's r | -0.75 | -0.51 | 0.66 |
|  | P-value | <0.001 | 0.06 | <0.001 |
| Subclinical<br>hyperthyroidism<br>(N = 175) | Slope (A) | -0.05±0.02 | -0.46±0.08 | 0.03±0.02 |
|  | Intercept (B) | -1.79±0.46 | 0.21±0.52 | 5.53±0.38 |
|  | Pearson's r | -0.13 | -0.38 | 0.10 |
|  | P-value | 0.07 | <0.001 | 0.17 |
| Overt<br>hyperthyroidism<br>(N = 75) | Slope (A) | 0.002±0.003 | 0.003±0.007 | 0.41±0.02 |
|  | Intercept (B) | -4.44±0.19 | -4.40±0.16 | -1.79±1.25 |
|  | Pearson's r | 0.07 | 0.04 | 0.91 |
|  | P-value | 0.56 | 0.72 | <0.001 |

**Fig. 1.**
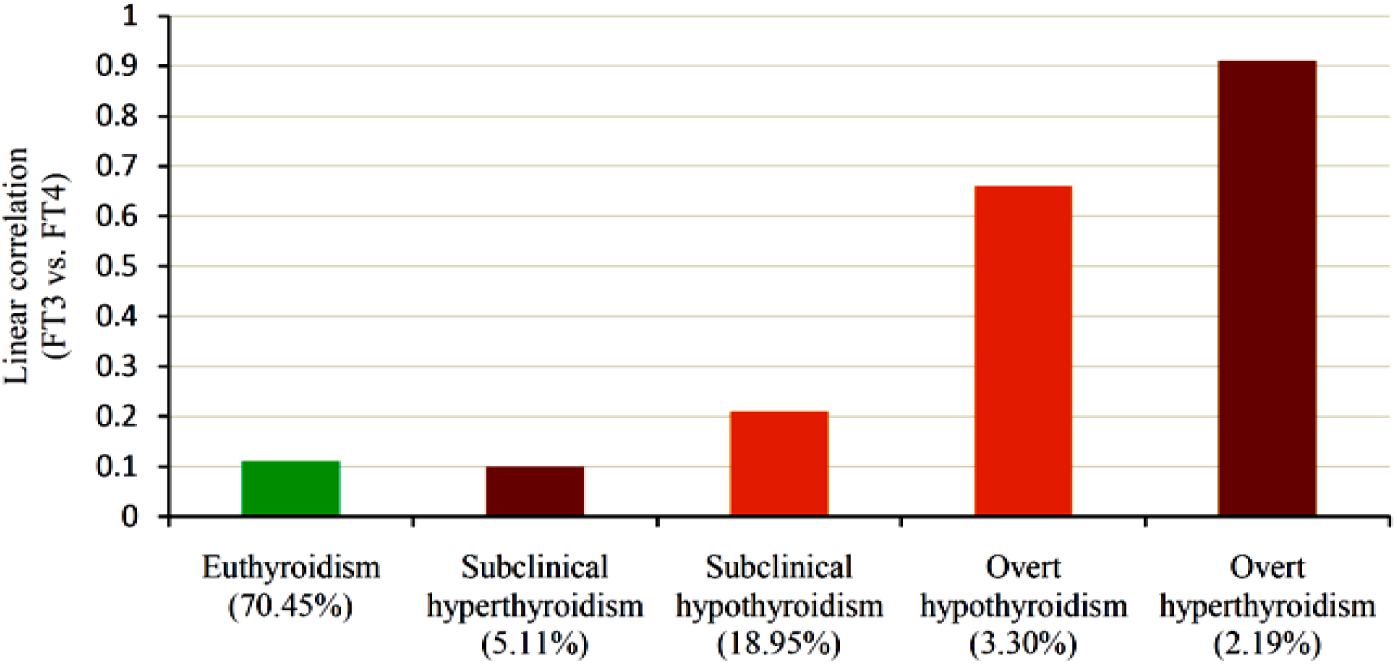
Increasing positive linear correlation between FT3 and FT4 from euthyroid to subclinical to overt thyroid dysfunctions where percentages in the brackets indicate the prevalence of thyroid disorders in 3425 (921M + 2504F) patients

The non-linear equation of ln(TSH) *vs* FT4 curve as a trend line between euthyroidism and hypothyroidism is given by the 4PL function (Eq. 2)

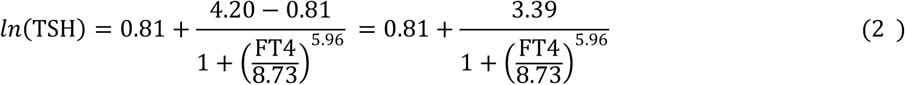

Comparing Eq. 2 with Eq. 1 and including standard errors in the coefficients, we observed the minimum asymptote (A) = 0.81±0.08 responding maximum FT4 = 28.20 pmol/L, the maximum asymptote (B) = 4.20±0.12 responding minimum FT4 = 0.88 pmol/L, inflection point (*x*_0_) = 8.73±0.30 pmol/L and Hill’s slope (*m*) = 5.96±1.07. This non-linear fit is symmetric with the nature of sigmoid curve.^2,28^ It has a stronger correlation (R^2^ = 0.97 and P <0.001). The second term in the right side of Eq. 2 appears from the effect of hypothyroidism.

The non-linear curve (Fig. 2-a) interestingly passes through the junction J of euthyroid, subclinical hypothyroid and overt hypothyroid regions. The point J can be taken as the turning point of normal to abnormal thyroid function and vice-versa. The patients suffering from overt hypothyroidism should have the history of changing TSH and FT4 through the curve C_1_JC_2_. Almost constant TSH along AC_1_ represents clinical euthyroidism, the curvature C_1_J arises due to subclinical hypothyroidism, JC_2_ demonstrates the biochemically degrading thyroid towards overt hypothyroidism and C_2_B shows FT4 falling to zero under the action of highly elevated TSH ≥100.00 mIU/L.

**Fig. 2.**
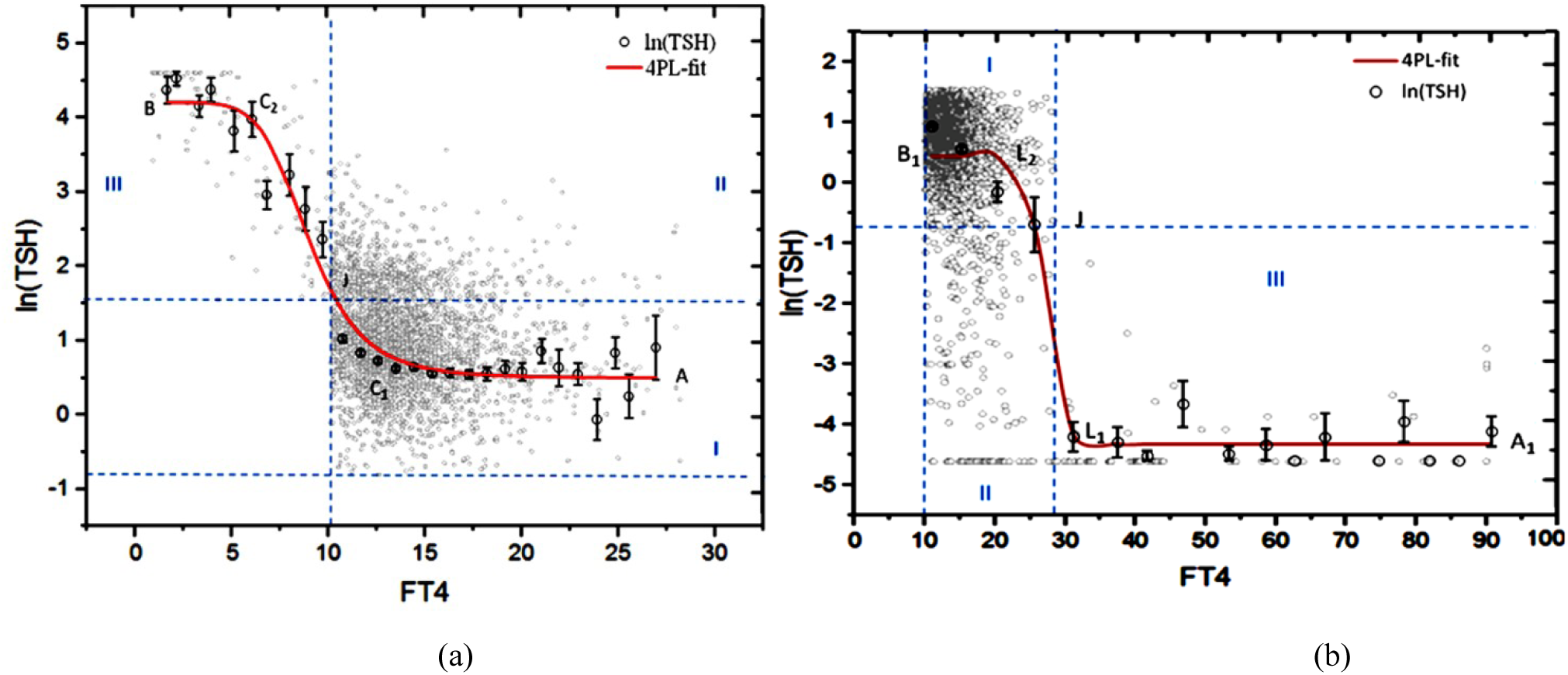
Non-linear relationships between TSH (mIU/L) and FT4 (pmol/L) governed by 4PL functions as the trend lines (R^2^ = 0.97 and p <0.001) between (a) euthyroidism-I and hypothyroidism-II to III regions, (b) euthyroidism-I and hyperthyroidism-II to III regions where the plots are drawn from the mean values related to FT4 bin size 1 pmol/L for (a) and 5 pmol/L for (b), the error bars indicate standard errors in ln(TSH), and number of subjects are 3175 (858M+2317F) in (a) and 2663 (728M+1935F) in (b).

In the Fig. 2-b, the 4PL curve of ln(TSH) *vs* FT4 gets steepness from euthyroid to subclinical hyperthyroid and to overt hyperthyroid regions of the data points. Stronger correlation given by the 4PL curve (R^2^ = 0.97, P <0.001) emphasizes for the non-linear relationship between TSH and FT4 in the patients suffering from euthyroidism to hyperthyroidism. In this 4PL function (Eq. 3), we find the minimum asymptote (A_1_) = -4.35±0.09 responding maximum FT4 = 90.00 pmol/L, the maximum asymptote (B_1_) = 0.20±0.19 responding minimum FT4 = 10.10 pmol/L, inflection point (*x*_0_) = 28.15±0.76 pmol/L and Hill’s slope (*m*) = 24.55±12.60.

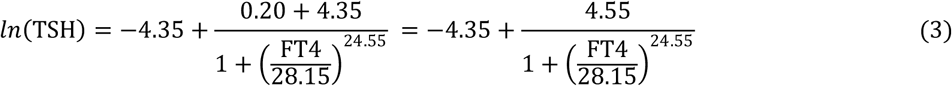

Referring to the Fig. 2-b, the hyperthyroidism develops in the patients from euthyroidism through the curve B_1_L_2_L_1_A_1_. Here, the linear fit passes through the junction J of euthyroid, subclinical and overt hyperthyroid regions whereas the 4PL curve passes through the subclinical hyperthyroid region. The steepness or Hill’s slope arises due to the negative correlation between TSH and FT4. The curvatures at points L_1_ and L_2_ come from the influence of overt hyperthyroidism and subclinical hyperthyroidism in the patients, respectively. The nearly straight line L_1_A_1_ signifies the continuous increment of FT4 up to 90.00 pmol/L at the lowest level of TSH ≤0.01 mIU/L in the overt hyperthyroid region.

It can be predicted that the patients under proper medication of overt hypothyroidism slowly return back to the normal condition through the curve BC_2_C_1_ (Fig. 2-a). In this way, the patients under proper medication of overt hyperthyroidism slowly return back to the normal condition through the curve A_1_L_1_L_2_ (Fig. 2-b). Finally, the gap: B-A = 3.39 in hypothyroidism (Eq. 2) totally vanishes when such thyroid disorder is completely cured. The non-zero second term in the right side of Eq. 1 refers to the hypothyroid dysfunction depending on the parameters: A, B, FT4, *x*_0_ and *m*. Obviously, A equals to B when Hill’s slope is *m* = 0. The minimum asymptote A refers to the normal TSH so that the maximum asymptote B should be corrected to A by reducing TSH for the subjects suffering from hypothyroidism. Similarly, the maximum asymptote B_1_ refers to the normal TSH so that the minimum asymptote A_1_ should be corrected to B_1_ by elevating TSH for the subjects suffering from hyperthyroidism.

The 4PL regressions establish the non-linear relationship between TSH and FT4 separately on hypothyroidism and hyperthyroidism interfaced with euthyroidism. The 4PL curve is better fitted with the observed data points giving rise to the biological interpretation of TSH response by FT4 deviation than other models such as linear, log-linear, non-linear sigmoid, error function and fourth-order polynomial.^1,2,8-^12, 28^^

Hypothyroidism and hyperthyroidism make the patients adaptive challenge to regulate the homeostatic system and to restore euthyroidism or ameliorate thyroid dysfunction. The effect of different functional states of the feedback system and the physiological variables such as ethnicity, age, iodine supply, T3 responsive features of nuclear receptors and many others add complexity to understand the functional relation between TSH and FT4. Here, ln(TSH) *vs* FT4 trend line differs in its nature as it passes from euthyroid to hypothyroid or to hyperthyroid set points. On the other hand, hysteresis in the HPT axis represents adaptive endocrine response by the evolutionarily conserved survival advantages.^29^ Temporarily persistent elevation of TSH despite the normalized FT4 levels indicates the transient hysteresis. Thyrotoxicosis is a state that leads to a lag time in recovery of TSH and a delayed recovery of T3 regulated genes. During the transition from thyrotoxicosis to euthyroidism, about 10% genes show incomplete recovery.^29^ In the recovery period of extreme hypo- and hyperthyroidism, the nature of ln(TSH) *vs* FT4 curves change showing the existence of hysteresis of HPT-axis. These facts are further associated with the different values of inflection point (*x*_0_) and Hill’s slope (*m*) resulting logistic curves governed by the Eq. 2 and Eq. 3 which are fitted graphically in the Fig. 2-a & 2-b for hypothyroidism and hyperthyroidism interfaced with euthyroidism. Berberich *et al*. (2018) performed a sensitivity analysis of a derived model revealing the dependency of TSH and thyroid hormones on the different system parameters by incorporating TSH-FT3-shunt inside the thyroid.^30^ Berberich’s study claims the understanding of thyroid hormone homeostasis developing a mathematical model of the HPT-loop. In our analysis, the deviation of ln(TSH) *vs* FT4 trend lines at the set points between euthyroidism and hypo/hyperthyroidism reflect the actual insight of the negative feedback mechanism in the thyroid cycle. T3, T4 and TSH have interlocking roles towards thyroid expressions, metabolic activities and regulations so that there exist complex interrelations among these variables.

## 4. Conclusion

In this study, the TFT analysis concludes that prevalence of hypothyroidism (18.95% subclinical and 3.30% overt hypothyroid patients) is higher than hyperthyroidism (5.11% subclinical and 2.19% overt hyperthyroid patients) in which females are more vulnerable than males and the thyroid abnormality increases with aging. The linear correlation between FT3 and FT4 becomes stronger in overt thyroid dysfunctions (0.66≤ r ≤0.91, P <0.001) than in subclinical hypo/hyperthyroidism and euthyroidism. This is important to understand the enzymatic action of monodeiodination converting T4 to T3 under the conditions of deregulated TSH. Most of the log-linear fits between TSH and FT4 (or FT3) related to these groups are statistically insignificant (0.001 ≤ P ≤ 0.72 and -0.03 ≤ r ≤ -0.75). The non-linear relation between ln(TSH) and FT4 is supported by the 4PL model (R^2^ = 0.97 and P <0.001) in both hypothyroidism and hyperthyroidism interfaced with euthyroidism. The curvatures in the 4PL curves are associated with the turning points of subclinical and overt thyroid dysfunctions from normal thyroid conditions and vice-versa. These curves are taken as the response trajectories of pituitary TSH by the deviation in serum FT4. The negative correlations and negative slopes in the log-linear plots are the indicators of negative feedback mechanism in the thyroid cycle.

## Ethical consent and Approval

This research work has been performed under the ethical guidelines provided by and taking authority from Institutional Review Board (IRB), Institute of Medicine (IOM), Tribhuvan University Teaching Hospital (TUTH), Maharajgunj, Kathmandu, Nepal.

## Acknowledgements

We are thankful to the head and team members of Department of Radiology, Department of Biochemistry and Institutional Review Board -TU Teaching Hospital for their technical supports in this research work.

## Data Availability

The data used to support the findings of this study are available from the corresponding author upon request.

